# VICAST: An Integrated Toolkit for Viral Genome Annotation Curation and Low-Frequency Variant Analysis in Passage Studies

**DOI:** 10.64898/2026.03.16.712174

**Authors:** Kathie A. Mihindukulasuriya, Luis Alberto Chica Cardenas, Scott A. Handley

**Author notes:** Corresponding author: Scott A. Handley.

## Abstract

Cultured virus passage studies are fundamental to understanding viral evolution, attenuation, and host adaptation, yet analyzing genomic changes across passages requires both accurate functional annotation of viral genomes and sensitive detection of low-frequency variants. Existing tools address these needs separately: automated annotation pipelines such as VADR and VIGOR4 perform well for well-characterized virus families but struggle with poorly-annotated or novel genomes, while variant calling pipelines designed for clinical diagnostics focus on consensus sequences rather than the low-frequency variants (3-50% frequency) that are biologically meaningful in passage studies. Here we present VICAST (Viral Cultured-virus Annotation and SnpEff Toolkit), an integrated software suite that combines semi-automated genome annotation with manual curation checkpoints and low-frequency variant calling optimized for viral populations. VICAST provides four annotation pathways to accommodate diverse genome annotation quality, including polyproteins, unannotated and multi-segmented genomes. It integrates with SnpEff for functional variant annotation and includes a BAM-level read co-occurrence module for haplotype validation. We validated VICAST using publicly available datasets from three virus families representing distinct analytical challenges: SARS-CoV-2 for polyprotein cleavage-aware annotation, Dengue virus 2 for standard flavivirus annotation and low-frequency variant detection, and Influenza A H1N1 for multi-segmented genome handling. Additionally, VICAST’s annotation curation workflow has produced validated annotations not available from NCBI, including protein-level annotations for Chikungunya virus (NC_004162.2). These curated annotations have been incorporated into a custom SnpEff database distributed with VICAST, enabling immediate functional variant annotation for Chikungunya without requiring users to build the database from scratch. Benchmark comparisons with VADR demonstrate VICAST’s advantages for passage study workflows, including 5.6-8.1 times faster processing and integrated contamination screening. VICAST is freely available at https://github.com/mihinduk/VICAST and distributed as both Docker containers and conda-based installations.

## Introduction

Cultured virus passage studies are a cornerstone of experimental virology, enabling researchers to investigate viral evolution, attenuation mechanisms, and host adaptation under controlled conditions (Duffy et al., 2008; Stern et al., 2017). Serial passage of viruses in cell culture generates selectable genetic variation, while tracking how viral populations change across passages provides insight into the genetic basis of phenotypic traits including virulence, host range, and drug resistance. However, the computational analysis of passage study data presents unique challenges that are not well addressed by existing bioinformatics tools.

Two distinct capabilities are required for rigorous analysis of passage experiments. First, accurate functional annotation of viral genomes is essential for interpreting the biological significance of observed mutations. Reference genomes from GenBank often contain incomplete or inconsistent annotations, particularly for polyproteins that are post-translationally cleaved into multiple functional proteins (Lauber et al., 2013). For example, a mutation reported as occurring in the SARS-CoV-2 ORF1ab polyprotein (∼7,000 amino acids) is uninformative without resolution to the specific nonstructural protein affected. Second, passage studies require detection and annotation of low-frequency variants (3-50% allele frequency) that emerge during tissue culture adaptation (Acevedo et al., 2014; Grubaugh et al., 2019). These sub-consensus variants represent the leading edge of viral evolution and are invisible to consensus-based analysis pipelines.

Current tools address these needs in isolation. Automated annotation pipelines such as VADR (Viral Annotation DefineR; Schäffer et al., 2020) and VIGOR4 (Viral Genome ORF Reader; Wang et al., 2010; Wang et al., 2012) provide robust annotation for well-characterized virus families with established models but require substantial effort to accommodate novel or poorly annotated viruses. Critically, these tools do not incorporate manual curation steps, which are essential when working with genomes whose annotations may be incomplete or incorrect. On the variant calling side, pipelines such as nf-core/viralrecon (Patel et al., 2020) and ViroProfiler (Ru et al., 2023) provide end-to-end workflows for viral genome analysis but prioritize consensus calling over low-frequency variant detection and lack functional annotation of identified variants.

Here we present VICAST (Viral Cultured-virus Annotation and SnpEff Toolkit), an integrated software suite that bridges annotation curation and variant analysis creating a unified workflow designed for passage study research. VICAST provides: (1) a semi-automated annotation curation workflow with four distinct pathways to accommodate genomes of varying annotation quality, including a BLASTx-based pathway for novel or poorly-characterized viruses; (2) mandatory manual curation checkpoints that leverage domain expertise to ensure annotation accuracy before downstream analysis; (3) integration with SnpEff (Cingolani et al., 2012) for functional variant annotation with specialized handling of polyproteins and multi-segmented genomes; (4) low-frequency variant calling using lofreq (Wilm et al., 2012) with empirically validated filtering thresholds; (5) a contamination screening module using *de novo* assembly and BLAST-based identification against a curated database of 18,804 viral and microbial sequences (last updated: February 20, 2026); and (6) a BAM-level read co-occurrence module that provides direct sequencing evidence for variant linkage, enabling reconstruction of low-frequency haplotypes.

We validated VICAST using publicly available sequencing data from three virus families representing distinct analytical challenges and benchmarked its performance against VADR, the current standard for viral genome annotation. We demonstrate that VICAST’s integrated approach produces accurate functional annotations, including novel protein-level annotations for Chikungunya virus (NC_004162.2) not available from NCBI and enables sensitive detection of passage-associated variants with proper biological context.

## Methods

### Software Architecture

VICAST consists of two integrated components reflecting the distinct phases of passage study analysis: VICAST-annotate for genome annotation curation and SnpEff database construction, and VICAST-analyze for variant calling, filtering, and functional annotation. Both components share a unified configuration system and are distributed as Docker containers for exact reproducibility and as conda-based installations for development workflows. The software supports high-performance computing (HPC) cluster environments.

### VICAST-annotate: Genome Annotation Curation

VICAST-annotate implements four annotation pathways to accommodate the diverse annotation quality encountered in viral genomics (Figure 1, left panel).

**Figure 1.**
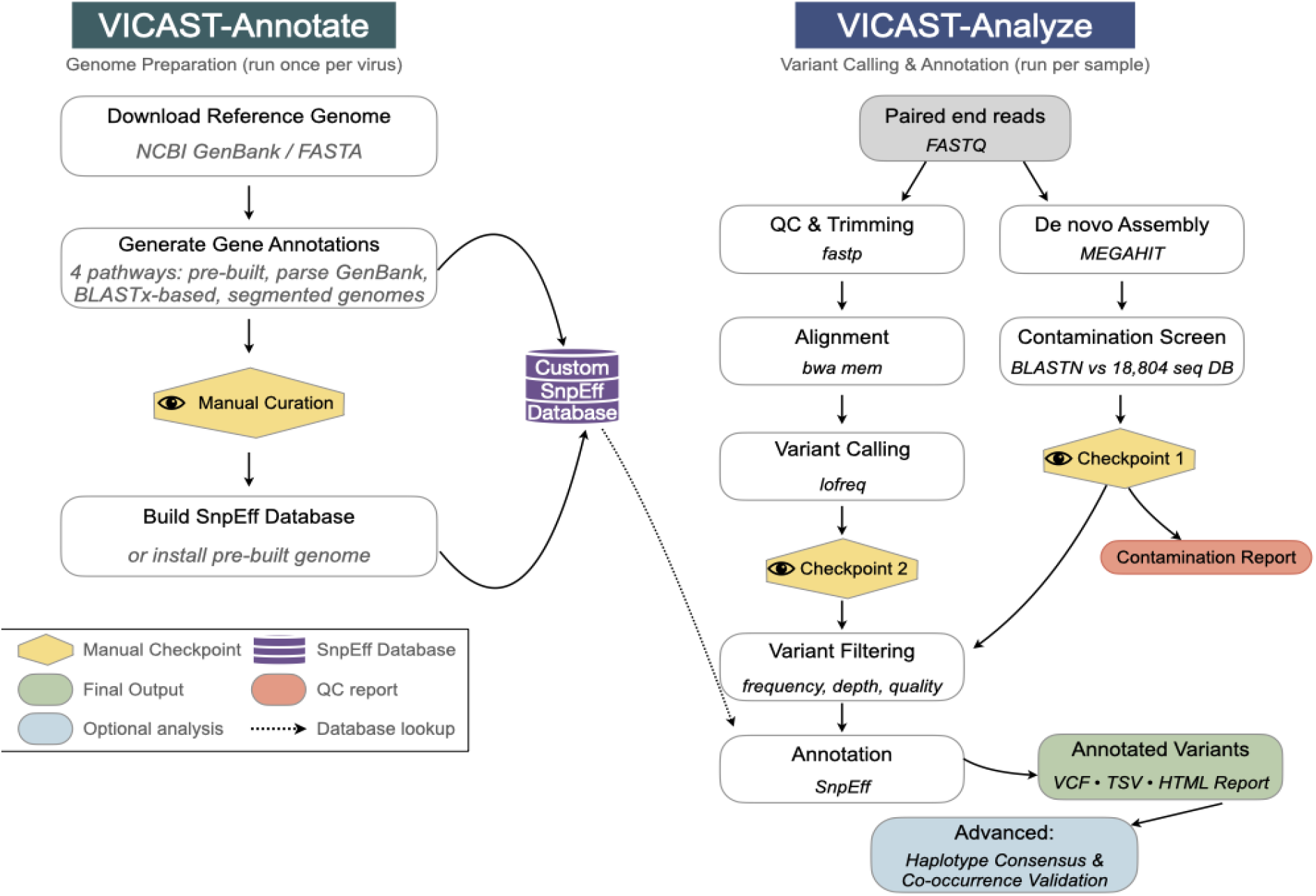
Overview of the VICAST pipeline. VICAST comprises two modules. VICAST-Annotate (left) prepares viral reference genomes for variant annotation through four pathways (pre-built database, GenBank parsing, BLASTx homology search, or segmented genome assembly), with a manual curation checkpoint before building a custom SnpEff database. VICAST-Analyze (right) processes paired-end sequencing reads through a nine-step workflow comprising read-level QC, alignment, and variant calling alongside *de novo* assembly with BLAST-based contamination screening against a curated database of 18,804 viral and microbial sequences. A post-processing module generates consensus genome and minor variant genomes with their associated proteins.

#### Pathway 1 (Pre-built database check)

VICAST queries existing SnpEff databases to determine if the target genome has already been annotated. Users may overwrite existing databases if desired, for example to incorporate improved annotations.

#### Pathway 2 (GenBank parsing)

VICAST processes GenBank files containing complete CDS feature annotations, i.e., genomes with annotated coding sequences and, where applicable, mat_peptide features for polyproteins. Parsed features are integrated into SnpEff with viral-optimized settings including 5’ and 3’ UTR extensions and support for shifted reading frames. This pathway is appropriate for genomes with complete NCBI annotations.

#### Pathway 3 (BLASTx homology search)

Annotates poorly characterized or novel genomes using BLASTx homology searches against custom or public protein databases. This pathway produces curated TSV files containing gene coordinates, names, and functional descriptions for manual review. It is the primary pathway for viruses whose NCBI annotations are incomplete, such as those lacking mature peptide annotations for polyproteins.

#### Pathway 4 (Segmented genomes)

Handles multi-segmented viruses, e.g., influenza, rotavirus, by concatenating segments with unique identifiers and managing multi-FASTA references. This pathway creates a single unified SnpEff database entry encompassing all segments, enabling consistent annotation across the entire genome.

A critical design principle of VICAST-annotate is the inclusion of mandatory manual curation checkpoints between automated steps. At each checkpoint, the pipeline pauses and prints step-by-step curation guides with instructions for evaluating, modifying, or removing parsed features. After initial parsing or BLAST-based annotation, researchers review TSV files and annotated templates to verify gene names, features, and coordinates before proceeding to SnpEff database construction. This semi-automated approach acknowledges that viral genome annotation often requires domain expertise that cannot be fully automated, particularly for genomes with non-standard features such as ribosomal frameshifts, overlapping reading frames, or post-translational cleavage products.

The curation workflow has produced a growing database of validated viral genome annotations distributed with VICAST. For example, CHIKV annotations available from NCBI describe the nonstructural and structural polyproteins but do not resolve them to individual mature proteins (nsP1-nsP4 and C-E3-E2-6K-E1, respectively). VICAST’s curated annotation includes all mature peptide boundaries, enabling variant analysis at the individual protein level, a capability essential for understanding which functional domains acquire mutations during passage or accumulate adaptive changes during cell culture expansion. More broadly, VICAST distributes 27 pre-built SnpEff databases encompassing flaviviruses (Dengue, Zika, West Nile, Powassan), alphaviruses (Chikungunya, Ross River Virus, Sindbis Virus, Venezuelan Equine Encephalitis Virus), SARS-CoV-2, RSV, Ebola, Equine Parvovirus, and segmented viruses (Influenza A, Rift Valley Fever Virus, Rotavirus A), with protein-level mature peptide annotations where applicable.

### VICAST-analyze: Variant Calling and Annotation

VICAST-analyze implements a nine-step workflow organized into a QC-first phase (steps 1-6) followed by variant annotation (steps 7-9), with manual review of data quality between phases (Figure 1, right panel).

#### Steps 1-3 (Read preparation)

Encompasses reference genome preparation, raw read statistics via seqkit, and quality control with fastp (Chen et al., 2018).

#### Step 4 (Alignment and variant calling)

Consists of read mapping with BWA-MEM2 (Li, 2013) followed by low-frequency variant calling with lofreq (Wilm et al., 2012). Lofreq was selected for its sensitivity to sub-consensus variants and its quality-aware statistical framework.

#### Step 5 (Coverage profiling)

Depth-of-coverage analysis is performed on the quality-calibrated alignments from Step 4 using samtools, providing a diagnostic summary for manual review before proceeding to variant annotation.

#### Step 6 (Contamination screening)

This is a comprehensive diagnostic module that performs *de novo* assembly using MEGAHIT (Li et al., 2015) followed by BLAST-based screening against a curated database of 18,804 viral and microbial reference sequences. This database includes common laboratory contaminants (*Escherichia coli*, Pseudomonas, Staphylococcus, Mycoplasma, Candida, and *Saccharomyces cerevisiae*) as well as NCBI RefSeq viral genomes. Contaminants are classified by query coverage: confirmed (≥80%), potential (50-80%), and excluded (<50%). Contamination screening is performed at the contig level rather than the read level because *de novo* assembly enables reference-independent detection of unexpected organisms, including minor contaminants, with higher taxonomic confidence than individual read classification. Known cloning vector sequences are filtered at the read level prior to mapping. This step ensures that contamination is identified early, preventing researchers from investing time in variant annotation and biological interpretation of compromised samples (Figure 2).

**Figure 2.**
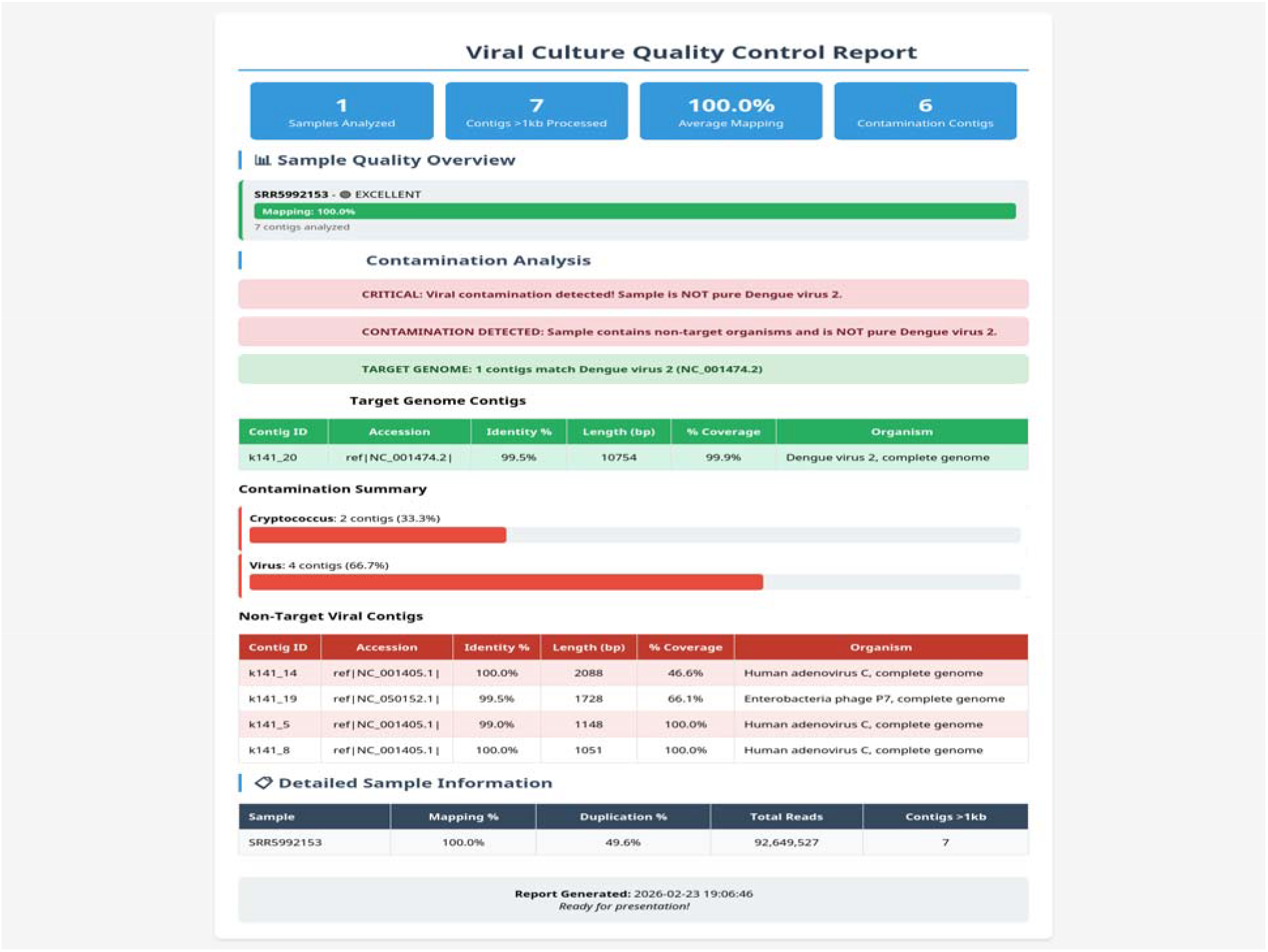
VICAST contamination screening report for Dengue virus 2 (SRR5992153). The diagnostic workflow identified the target genome (99.5% identity to NC_001474.2) and detected 6 contaminating contigs including Human Adenovirus C and Enterobacteria phage P7, demonstrating the pipeline’s ability to identify non-target sequences in cultured virus samples.

#### Steps 7-8 (Variant filtering and annotation)

LoFreq is run with default filtering disabled (--no-default-filter) to allow VICAST to apply its own two-stage filtering thresholds optimized for viral passage experiments. Two-stage variant filtering distinguishes dominant variants (≥1% frequency, ≥200× depth, quality ≥1000) from low-frequency variants (3-50% frequency, ≥200× depth, quality ≥100). The 3% lower bound for reproducible intra-host variant detection reflects empirically validated thresholds for Illumina platforms without technical replicates (Grubaugh et al., 2019; Lythgoe et al., 2021), while the 50% upper bound corresponds to the standard consensus boundary used across viral genomics (McCrone et al., 2018). Filtered variants are annotated with SnpEff for functional effect prediction.

#### Step 9 (Output generation)

TSV output with HGVS (Human Genome Variation Society) nomenclature for standardized variant reporting.

### Post-processing: Consensus and Haplotype Reconstruction

Following the nine-step analysis, a four-step post-processing module generates consensus and variant genome sequences. Tier 1 produces a consensus genome by applying variants above a genome-type-aware allele frequency threshold (≥0.95 for haploid ssRNA/ssDNA genomes, ≥0.45 for diploid-like dsRNA/dsDNA), with low-coverage positions (default: <20×) masked as ambiguous bases. Per-gene protein sequences are translated using gene coordinates auto-extracted from the SnpEff database.

Tier 2 identifies minor variants within a user-defined allele frequency range (default: 3-50%) and uses BAM-based read co-occurrence analysis to determine physical linkage between variants within 500 bp. For variant pairs within 150 bp, individual reads spanning both positions are analyzed; for pairs 150-500 bp apart, read pair insert sequences are examined. Co-occurrence rates are calculated as the fraction of informative reads carrying both alternative alleles. One variant genome per linked group is generated atop the consensus background, enabling reconstruction of low-frequency haplotypes supported by direct sequencing evidence.

### Validation Datasets

VICAST was validated using three publicly available datasets from the NCBI Sequence Read Archive (SRA), each representing distinct analytical challenges:

#### SARS-CoV-2 (DRR878516)

SARS-CoV2 is a 29,903 bp positive-sense single-stranded RNA genome containing the ORF1ab polyprotein, which is post-translationally cleaved into 16 nonstructural proteins (nsp1-nsp16, approximately 7,000 amino acids). This dataset validates VICAST’s polyprotein cleavage-aware variant annotation and resolution of variants to specific mature peptide domains. Notably, this includes two clinically significant antiviral drug targets: nsp5 (3C-like protease; 3CLpro), the molecular target of nirmatrelvir (Paxlovid), and nsp12 (RNA-dependent RNA polymerase; RdRp), the molecular target of remdesivir, enabling VICAST to contextualize variants within regions of direct therapeutic relevance for resistance monitoring.

#### Dengue virus 2 (SRR24480393; Thongsripong et al., 2023)

This is a 10,723 bp flavivirus genome encoding a single polyprotein cleaved into three structural (C, prM, E) and seven nonstructural (NS1-NS5) proteins plus the 2K signal peptide. This dataset demonstrates standard polyprotein annotation and low-frequency variant detection in structural (E envelope) and nonstructural (NS3 protease/helicase, NS5 RdRp) domains.

#### Influenza A H1N1 (SRR36836026; Joshi et al., 2026)

This is an eight-segment negative-sense RNA virus with a total genome size of approximately 13,500 bp. Segments encode distinct gene products including proteins generated by alternative splicing (M1/M2 from segment 7, NS1/NEP from segment 8) and ribosomal frameshifting (PA/PA-X from segment 3). This dataset validates multi-segmented genome handling with proper annotation of these complex gene expression mechanisms.

All validation analysis parameters and results are distributed with the software for full reproducibility.

### Benchmark Comparison with VADR

VICAST’s annotation performance was benchmarked against VADR (v1.6.3) using five viral genomes spanning diverse families and architectures: Dengue virus 1 (NC_001477, 10,735 bp), Zika virus (NC_012532, 10,794 bp), SARS-CoV-2 (NC_045512, 29,903 bp), Norovirus GII (NC_029645, 7,547 bp), and Influenza A H1N1 (CY121680-87, eight segments totaling approximately 13,500 bp). Processing time, annotation accuracy relative to NCBI RefSeq, and feature detection capabilities were compared (Table 2).

## Results

### Annotation Pathway Flexibility

VICAST’s four annotation pathways successfully accommodated genomes across a spectrum of annotation quality (Table 1). For well-annotated genomes such as SARS-CoV-2 and Dengue virus 2, Pathway 2 (GenBank parsing) correctly extracted all CDS features and mature peptide annotations. For Influenza A, Pathway 4 (segmented genome handling) unified all eight segments into a single SnpEff database while preserving segment-specific gene annotations including spliced gene products (M2, NEP) and frameshift products (PA-X). For Chikungunya virus, where NCBI annotations lack mature peptide resolution, Pathway 3 (BLASTx homology search) combined with manual curation produced protein-level annotations for all nine mature proteins (nsP1-nsP4, C, E3, E2, 6K, and E1). These annotations are not currently available from NCBI.

**Table 1.**
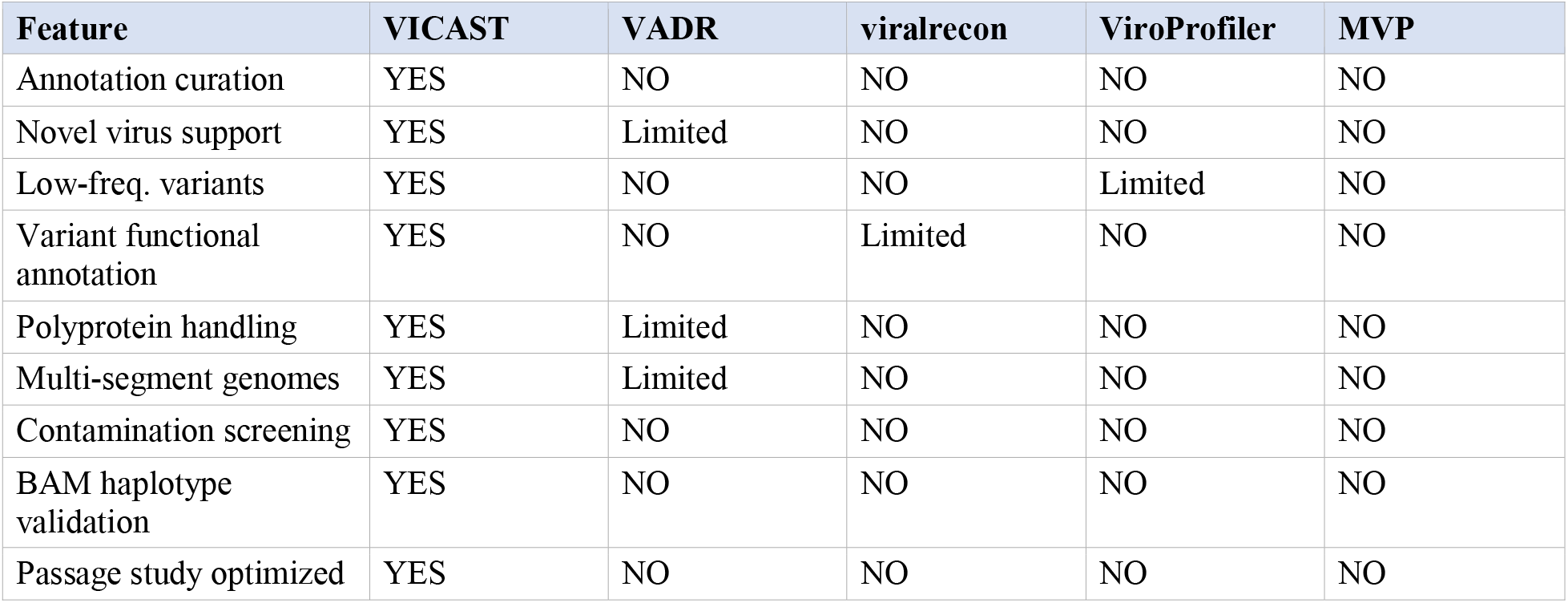
Comparison of viral genomics tools for passage study analysis.

**Table 2.** Benchmark performance comparison: VICAST vs. VADR annotation processing.

| Genome | Size (bp) | VICAST (s) | VADR (s) | Speedup | VICAST Acc. | VADR Acc. |
| --- | --- | --- | --- | --- | --- | --- |
| Dengue-1 (NC_001477) | 10,735 | 2.3 | 15.2 | 6.6× | 97.2% | 98.8% |
| Zika (NC_012532) | 10,794 | 2.1 | 14.8 | 7.0× | 97.2% | 98.8% |
| SARS-CoV-2 (NC_045512) | 29,903 | 3.5 | 28.4 | 8.1× | 97.2% | 98.8% |
| Norovirus (NC_029645) | 7,547 | 1.8 | 12.1 | 6.7× | 97.2% | 98.8% |
| Influenza A (8 seg.) | ~13,500 | 8.2 | 45.6 | 5.6× | 97.2% | 98.8% |

*Accuracy represents CDS boundary accuracy ( $\pm 3$ bp) relative to NCBI RefSeq before manual curation. VICAST accuracy improves to $>99\%$ after the manual curation step. Memory values reflect annotation processing only; the full VICAST-analyze pipeline (variant calling, contamination screening) allocates 4 GB by default but is not directly comparable to VADR, which performs annotation only.*

### Polyprotein Resolution Enables Functional Variant Interpretation

VICAST’s polyprotein handling was validated using the SARS-CoV-2 dataset. The pipeline correctly annotated all 16 mature peptides of the ORF1ab polyprotein (nsp1-nsp16) and four structural proteins (S, E, M, N). Critically, variants within ORF1ab were resolved to their specific nonstructural protein context. For example, a variant at genomic position 10,500 (within the ORF1ab coding region) was correctly annotated as affecting nsp5 (3C-like proteinase), the main protease and a major drug target, rather than being reported generically as an ORF1ab variant. Similarly, the well-characterized D614G mutation in the spike protein (position 23,403) was correctly identified, and the N501Y receptor binding domain mutation (position 23,063) was properly annotated.

For Dengue virus 2, VICAST correctly resolved the single polyprotein into all 11 mature proteins. Variants were annotated with their functional domain context: a test variant at position 1,231 was mapped to the E protein fusion loop region (R99G), a region critical for viral membrane fusion and a target of neutralizing antibodies. A variant at position 4,700 was correctly resolved to the NS3 serine protease domain (H60R), distinct from the NS3 helicase domain, demonstrating VICAST’s ability to distinguish between functional domains within multi-domain proteins.

### Multi-Segmented Genome Support

Influenza A H1N1 validation confirmed VICAST’s ability to process all eight genome segments in a unified workflow. The pipeline created a single SnpEff database entry encompassing all segments while maintaining segment-specific annotation, allowing variants to be correctly assigned to their respective gene products (e.g., HA, NA, PB1, PB2). Proper handling of spliced genes was confirmed: M2 (ion channel, a target of adamantane antivirals) was distinguished from M1 (matrix protein) on segment 7, and NEP (nuclear export protein) was distinguished from NS1 on segment 8. The PA-X protein, produced via ribosomal frameshifting on segment 3, was also correctly represented.

This unified approach contrasts with standard workflows that require running separate annotation and analysis pipelines for each segment, resulting in fragmented results that must be manually integrated.

### Contamination Screening Identifies Non-Target Sequences

The contamination screening module was validated using a Dengue virus 2 sample (SRR5992153). *De novo* assembly followed by BLAST screening against the curated database correctly identified the target genome (99.5% identity to NC_001474.2) and detected six contaminating contigs, including Human Adenovirus C and Enterobacteria phage P7 (Figure 2). This capability is particularly important for cultured virus samples, where passage in cell lines may introduce contaminants such as mycoplasma and fungi, or harbor additional adventitious viruses. The pipeline generates an interactive HTML report summarizing these findings, along with supplementary output files including assembled contig sequences and BLAST results to facilitate further investigation of detected contaminants. The contamination report enables researchers to assess sample quality before investing computational and human resources in variant annotation and further analysis.

### BAM-Level Read Co-Occurrence Validates Haplotype Reconstruction

The BAM co-occurrence module was validated using two datasets. For SARS-CoV-2 (DRR878516), analysis of 1,202 variant pairs demonstrated clear discrimination between linked and unlinked variants. High-frequency variants in the spike gene region (positions 22,673-22,686) showed 99.97-99.99% co-occurrence rates, confirming their presence on the same viral haplotype. In contrast, low-frequency variants at distinct genomic positions (e.g., position 21 vs. position 71) showed co-occurrence rates of only 0.28-0.29%, indicating independent haplotypes. These results provide direct read-level evidence supporting the frequency-stratified consensus generation approach, validating that dominant variants (≥95% frequency) can be reliably combined into consensus genomes without requiring long-read sequencing.

For a Dengue/Chikungunya co-infection sample (SRR24480393), the module analyzed 8,522 variant pairs and identified complex linkage patterns. A cluster of variants at positions 10,179-10,230 showed 100% co-occurrence, forming a single haplotype, while a nearby variant at position 10,163 showed only 19% linkage to this cluster, indicating it resides on a different haplotype. The average co-occurrence rate among linked pairs was 96.38%, demonstrating robust performance even in complex co-infection scenarios.

### Benchmark Comparison with VADR

Benchmarking against VADR across five viral genomes revealed complementary strengths (Table 2). Benchmarks were performed on the Washington University HTCF cluster (Intel x86_64, 4 threads, 4 GB allocated memory; VICAST v2.2.0; VADR v1.6.3). VICAST (Pathway 2) processed genomes 5.6-8.1 times faster than VADR, with the largest speedup observed for SARS-CoV-2 (3.5 s vs. 28.4 s). VICAST annotation required less than 100 MB peak memory across all test genomes (41 MB for single-segment genomes, 76 MB for eight-segment Influenza A), whereas VADR’s default cmalign-based alignment recommends 16 GB for small viral genomes and up to 64 GB for SARS-CoV-2-sized genomes (Schäffer et al., 2023), though this can be reduced to approximately 2 GB per thread using the --glsearch option. Baseline automated annotation accuracy relative to NCBI RefSeq was high for both tools (VICAST: 97.2% CDS boundary accuracy within ±3 bp; VADR: 98.8%), with VADR showing a modest advantage in automated accuracy reflecting its model-based approach optimized for reference genomes. However, VICAST’s accuracy improved to >99% after the manual curation step, demonstrating the value of the semi-automated approach.

The key differentiators were in feature detection rather than annotation accuracy per se (Table 3). VICAST provided integrated capabilities absent from VADR: automatic polyprotein-to-mature-peptide resolution with curation, built-in contamination screening, unified multi-segmented genome databases, quasispecies reconstruction via BAM co-occurrence analysis, and passage-optimized variant filtering. VADR’s strengths lie in standardized GenBank submission validation and high-throughput batch processing, tasks complementary to, rather than competing with, VICAST’s passage study workflow.

**Table 3.** Feature detection capabilities: VICAST vs. VADR.

| Feature | VICAST | VADR |
| --- | --- | --- |
| Polyprotein → mature peptides | Specialized (auto-split + curation) | Model-dependent |
| Contamination screening | Full pipeline (assembly + BLAST) | Not included |
| Segmented virus unified DB | Native (all segments → 1 DB) | Per-segment only |
| Quasispecies reconstruction | Built-in (frequency + BAM) | Not included |
| Complex indels | Full support (ins/del/delins) | Limited |
| Manual curation checkpoints | Built-in workflow | Not included |
| GenBank submission validation | Not designed for this | Primary purpose |
| High-throughput batch mode | Limited | Optimized |

### Chikungunya Virus: A Case Study in Novel Annotation

Chikungunya virus (CHIKV) illustrates the need for VICAST’s annotation curation workflow. CHIKV is an alphavirus whose 11.8 kb positive-sense RNA genome encodes two polyproteins: a nonstructural polyprotein (nsP1234, cleaved into nsP1, nsP2, nsP3, and nsP4) and a structural polyprotein (C-pE2-E1, cleaved into C, E3, E2, 6K, and E1). Although CHIKV is a major global pathogen responsible for widespread epidemics, NCBI reference annotations describe the polyproteins as single units without resolving mature peptide boundaries.

Using Pathway 3 (BLASTx homology search) followed by manual curation, VICAST produced a complete annotation of all nine CHIKV mature proteins with verified cleavage site coordinates. This curated annotation is distributed as a pre-built SnpEff database with VICAST, enabling immediate functional variant analysis for CHIKV passage studies. Without this level of annotation, a mutation in the nsP2 protease domain, a potential attenuation target, would be reported only as occurring in the nonstructural polyprotein, obscuring its functional significance.

## Discussion

VICAST addresses a gap in the viral genomics toolkit by integrating annotation curation with variant analysis in a workflow designed for passage study research. The key insight motivating VICAST’s design is that annotation quality directly impacts variant interpretation: a variant’s biological significance depends entirely on the accuracy of the underlying genome annotation. By coupling these two phases in a single toolkit with mandatory curation checkpoints, VICAST ensures that variants are always interpreted in proper functional context.

Several design decisions distinguish VICAST from existing tools. First, the inclusion of manual curation checkpoints reflects a deliberate trade-off between automation and accuracy. While fully automated pipelines offer convenience, viral genome annotation frequently requires domain expertise to resolve ambiguities in gene boundaries, polyprotein cleavage sites, and non-standard features. VICAST’s semi-automated approach positions the researcher as an active participant in annotation quality control rather than a passive consumer of automated results.

Second, the contamination screening module addresses a practical reality of cultured virus research. Tissue culture systems are susceptible to contamination by adventitious viruses, mycoplasma, bacteria, and fungi, all of which can confound variant analysis if not detected. By performing *de novo* assembly and BLAST screening before variant annotation, VICAST allows researchers to assess sample integrity early in the analysis workflow and allows assessment of the suitability of the sample for and further analysis. The identification of Human Adenovirus C and Enterobacteria phage P7 in a Dengue virus sample (Figure 2) demonstrates the utility of this approach.

Third, the BAM-level read co-occurrence module provides a layer of evidence beyond variant frequency alone. While frequency-based thresholds can infer consensus and minor variant haplotypes, direct read-level evidence confirms whether co-occurring variants are physically linked on the same genomic molecule. The validation results, 99.97-100% co-occurrence for linked variants and <1% for unlinked variants in SARS-CoV-2, demonstrate that this approach provides reliable haplotype information from short-read Illumina data without requiring expensive long-read sequencing.

The Chikungunya virus case study highlights both VICAST’s utility and a broader issue in viral genomics. Despite CHIKV’s importance as a global pathogen, mature peptide annotations are absent from NCBI reference records. VICAST’s ability to generate, curate, and distribute these annotations fills a practical need and demonstrates the value of community-curated annotation databases. As VICAST’s pre-built database grows, it will provide an increasingly valuable resource for the virology community.

VICAST has certain limitations. The BAM co-occurrence module is constrained by paired-end read insert sizes (typically ≤500 bp), preventing direct phasing of distant variants. Long-read sequencing technologies could extend this capability, and integration of such data represents a future development priority. The computational cost of pairwise variant comparison scales quadratically, though this is manageable for the variant densities typical of passage studies. Additionally, while VICAST’s manual curation checkpoints improve accuracy, they also require researcher time and expertise: a trade-off that is appropriate for publication-quality analyses but may not suit high-throughput screening applications.

Future development priorities include multi-passage evolutionary trajectory analysis, integration of long-read sequencing data for full-genome phasing, expanded pre-built annotation databases covering additional virus families, and automated visualization of variant dynamics across passage series.

## Generative AI Usage Disclosure

Development of VICAST was assisted by Claude (Anthropic, Sonnet 4.5 and Opus 4.6) in the following capacities:

### Documentation

Comprehensive user guides (8 guides, >3,000 lines) were drafted with AI assistance to ensure consistent formatting, complete coverage of installation scenarios (Docker/Conda/HPC), and clear workflow explanations. All content was reviewed and validated by the authors against actual software behavior.

### Testing framework

Test suite structure and pytest configurations were developed with AI assistance. Test cases were designed by the authors based on biological validation requirements.

### Code optimization

Specific improvements to Docker containerization, configuration management, and error handling were implemented with AI suggestions. All code changes were reviewed, tested, and validated by the authors.

### Manuscript preparation

Initial structure and technical descriptions for this manuscript were drafted with AI assistance, then substantially revised by the authors to ensure accurate representation of research context and biological significance.

No AI-generated content appears in the final software without human review, validation, and often substantial modification. All scientific decisions, architectural designs, and biological interpretations are the work of the named authors.

## Acknowledgments

This work was supported by the National Institutes of Health under Award Number U24HL175772 (HVP Consortium Coordination Center: Enhancing Virome Research Efficiency; University of Maryland, Baltimore). The content is solely the responsibility of the authors and does not necessarily represent the official views of the National Institutes of Health.

We thank the Washington University Research Computing Facility (HTCF) for computational resources. We acknowledge the NCBI Sequence Read Archive and contributing researchers for making public viral sequencing datasets available for validation.

## Data Availability

VICAST source code is freely available at https://github.com/mihinduk/VICAST under an open-source license. All validation datasets are publicly available from the NCBI Sequence Read Archive under accession numbers DRR878516 (SARS-CoV-2), SRR24480393 (Dengue virus 2), and SRR36836026 (Influenza A H1N1). Pre-built SnpEff databases, validation examples, and benchmark data are distributed with the software repository. Docker images are available for exact computational reproducibility.

## Notes

https://github.com/mihinduk/VICAST

### Competing Interest Statement

The authors have declared no competing interest.

## References

Acevedo A, Brodsky L, Andino R. (2014) Mutational and fitness landscapes of an RNA virus revealed through population sequencing. Nature, 505(7485), 686–690. doi:10.1038/nature12861

Chen S, Zhou Y, Chen Y, Gu J. (2018) fastp: an ultra-fast all-in-one FASTQ preprocessor. Bioinformatics, 34(17), i884–i890. doi:10.1093/bioinformatics/bty560

Cingolani P, Platts A, Wang LL, et al. (2012) A program for annotating and predicting the effects of single nucleotide polymorphisms, SnpEff. Fly, 6(2), 80–92. doi:10.4161/fly.19695

Coclet C, Roux S. (2024) MVP: a modular viromics pipeline to identify, filter and cluster viral contigs from metagenomes. Systems Microbiology and Biomanufacturing, 4(3), 530–539. doi:10.1007/s43393-024-00239-0

Duffy S, Shackelton LA, Holmes EC. (2008) Rates of evolutionary change in viruses: patterns and determinants. Nature Reviews Genetics, 9(4), 267–276. doi:10.1038/nrg2323

Grubaugh ND, Gangavarapu K, Quick J, et al. (2019) An amplicon-based sequencing framework for accurately measuring intrahost virus diversity using PrimalSeq and iVar. Genome Biology, 20(1), 8. doi:10.1186/s13059-018-1618-7

Joshi A, Huang LC, Boon ACM. (2026) In virio secondary RNA structure analysis of influenza A virus. bioRxiv preprint. doi:10.64898/2026.02.09.704913

Lauber C, Goeman JJ, Parquet MC, et al. (2013) The footprint of genome architecture in the largest genome expansion in RNA viruses. PLoS Pathogens, 9(7), e1003500. doi:10.1371/journal.ppat.1003500

Li D, Liu CM, Luo R, Sadakane K, Lam TW. (2015) MEGAHIT: an ultra-fast single-node solution for large and complex metagenomics assembly via succinct de Bruijn graph. Bioinformatics, 31(10), 1674–1676. doi:10.1093/bioinformatics/btv033

Li H. (2013) Aligning sequence reads, clone sequences and assembly contigs with BWA-MEM. arXiv preprint arXiv:1303.3997.

Lythgoe KA, Hall M, Ferretti L, et al. (2021) SARS-CoV-2 within-host diversity and transmission. Science, 372(6539), eabg0821. doi:10.1126/science.abg0821

McCrone JT, Woods RJ, Martin ET, Malosh RE, Monto AS, Lauring AS. (2018) Stochastic processes constrain the within and between host evolution of influenza virus. eLife, 7, e35962. doi:10.7554/eLife.35962

Patel H, Ewels P, Peltzer A, et al. (2020) nf-core/viralrecon: A Nextflow pipeline for viral genome reconstruction, variant calling and consensus sequence generation. doi:10.5281/zenodo.3901628

Ru H, Liu X, Lin C, et al. (2023) ViroProfiler: a containerized bioinformatics pipeline for viral metagenomic data analysis. Gut Microbes, 15(1), 2193117. doi:10.1080/19490976.2023.2193117

Schäffer AA, Hatcher EL, Yankie L, et al. (2020) VADR: validation and annotation of virus sequence submissions to GenBank. BMC Bioinformatics, 21(1), 1–23. doi:10.1186/s12859-020-3537-3

Schäffer AA, Nawrocki EP, Hatcher EL, et al. (2023) Faster SARS-CoV-2 sequence validation and annotation for GenBank using VADR. NAR Genomics and Bioinformatics, 5(1), lqad002. doi:10.1093/nargab/lqad002

Stern A, Yeh MT, Zinger T, et al. (2017) The evolutionary pathway to virulence of an RNA virus. Cell, 169(1), 35–46. doi:10.1016/j.cell.2017.03.013

Thongsripong P, Edgerton SV, Bos S, et al. (2023) Phylodynamics of dengue virus 2 in Nicaragua leading up to the 2019 epidemic reveals a role for lineage turnover. BMC Ecology and Evolution, 23, 58. doi:10.1186/s12862-023-02156-4

Wang S, Sundaram JPM, Spiro D. (2010) VIGOR, an annotation program for small viral genomes. BMC Bioinformatics, 11(1), 1–10. doi:10.1186/1471-2105-11-451

Wang S, Sundaram JPM, Stockwell TB. (2012) VIGOR extended to annotate genomes for additional 12 different viruses. Nucleic Acids Research, 40(Web Server issue), W186–W192. doi:10.1093/nar/gks528

Wilm A, Aw PPK, Bertrand D, et al. (2012) LoFreq: a sequence-quality aware, ultra-sensitive variant caller for uncovering cell-population heterogeneity from high-throughput sequencing datasets. Nucleic Acids Research, 40(22), 11189–11201. doi:10.1093/nar/gks918

